# Organoid-based modeling of platinum resistance identifies KRT17 as both a response mediator and biomarker for targeted therapy in ovarian cancer

**DOI:** 10.1101/2025.03.10.642373

**Authors:** Juliane Reichenbach, Juliana Schmid, Sophia Hierlmayer, Tingyu Zhang, Ilaria Piga, Sophia Geweniger, Jonas Fischer, Aarushi Davesar, Nemanja Vasovic, Anca Chelariu-Raicu, Fabian Kraus, Alexander Burges, Bastian Czogalla, Doris Mayr, Tobias Straub, Christoph Klein, Jesper V. Olsen, Sven Mahner, Fabian Trillsch, Mirjana Kessler

## Abstract

Variable platinum responses drive high mortality in high-grade serous ovarian cancer (HGSOC), but to date, there is no molecular approach to define resistance levels for clinical decision-making. Here, we developed the organoid drug resistance assay (ODR-test) with patient-derived organoids from our ovarian cancer biobank and found that all HGSOC patients develop molecular resistance under exposure to carboplatin, although with varying clinical implications. Sustained phenotypic reprogramming and cellular plasticity under carboplatin pressure emerged as a conserved mechanism irrespective of the basal resistance level. Transcriptional and proteomic analyses revealed changes in cell adhesion and differentiation in post-platinum lines as adaptive responses that drive the increase in resistance. We identified Keratin 17 (KRT17) as a mediator of developing platinum resistance and validated its function by CRISPR/Cas9 and overexpression. Additionally, we found that KRT17 expression status (K-score) is a significant negative prognostic histopathological biomarker in a large cohort (N=384) of advanced HGSOC patients. In organoids, increased KRT17 levels enhanced sensitivity to PI3K/Akt inhibitors Alpelisib and Afuresertib, highlighting the potential of KRT17 as a stratification biomarker for targeted therapies.

## Introduction

The pronounced heterogeneity of high-grade serous ovarian cancer (HGSOC) challenges the development of targeted therapies, leaving platinum-based chemotherapy as the main backbone of treatment ^1^. However, while initially effective in the vast majority of patients, its therapeutic benefits diminish over time, as tumors acquire resistance, resulting in highly variable response rates among patients ^2^ ^3^. In recurrent disease, the decision regarding platinum re-administration still relies only on clinical and chronological criteria due to the absence of molecular predictive and stratification tools ^4^. Consequently, many patients receive ineffective therapy, and therapeutic options remain limited ^5^. Despite extensive research into the molecular adaptions that tumors undergo during platinum treatment, preclinical studies have not yet yielded clinically applicable results ^6^ ^7^. As an important advancement to standard cell lines and cell line-based xenograft models, patient-derived organoids (PDOs) of HGSOC maintain the epithelial structure, allow long-term cultivation, and preserve the full genomic and functional hallmarks of ovarian cancer ^8–11^. Thus, an HGSOC PDO biobank from a large heterogeneous prospective patient cohort with well-documented clinical data provides a unique resource for studying individual differences in platinum response and the emergence of resistant phenotypes ^12^.

Standard viability-based *in vitro* drug assays quantify only the direct cytotoxicity of chemotherapy by measuring the cell viability after exposure ^13^. However, since therapy in patients likely drives selection mechanisms within a tumor that allows the expansion of pre-existing minor clones, viability testing is inadequate for capturing this critical aspect of drug response ^14^ ^15^.

Understanding the role of changes in tumor stemness and cell biology over time is critical to unraveling the mechanisms of platinum resistance. Transcriptional profiling of the fallopian tube epithelium, the origin of HGSOC ^16^, revealed subclusters of cells with stem cell signatures ^17–20^. Among these, the type I intermediary cytoskeletal filament protein Keratin 17 (KRT17) has been identified as one of the markers of the progenitor population ^17–20^. In other epithelial tissues, KRT17 is studied for its role in tissue regeneration and stress responses ^21^ ^22^. By activating the PI3K/Akt signaling cascade, KRT17 promotes stemness and cell fate change in keratinocytes ^23^. Upregulated in various epithelial cancers, including HGSOC, KRT17 correlates with poor clinical outcomes, cell proliferation, and therapy resistance ^24–34^. Notably, inhibition of the PI3K/Akt signaling cascade has been proposed to target chemoresistance in ovarian cancer ^35–37^.

In this study, we have developed the *in vitro* Organoid Drug Resistance Test (ODR-test) to quantify the effect of chemotherapy, in this case platinum, on cancer stemness potential. Long-term cultivation of *post*-platinum-treated HGSOC PDOs enabled transcriptional and proteomic analyses of adaptive changes under therapeutic pressure. We identified KRT17 as a mediator of platinum response and validated our findings of a potential clinical prognostic impact in a large cohort of advanced HGSOC patients. Notably, high KRT17 levels could stratify the use of PI3K/Akt-inhibitors in HGOSC treatment, with organoids that overexpressed KRT17 showing improved sensitivity to the PI3K-inhibitor Alpelisib and Akt-inhibitor Afuresertib. These findings pave the way for the identification of platinum-resistant ovarian cancer using biomarkers and for individualized, molecular-based treatment strategies that improve the management following suboptimal responses to platinum chemotherapy.

## Results

### The Organoid-Drug-Resistance Test

Primary tumor tissue from HGSOC patients, obtained during ovarian cancer debulking surgeries was processed and patient-derived organoid (PDO) lines were generated as described previously within our ovarian cancer organoid biobank ^12^. To investigate patient-specific platinum responses, we selected four organoid lines (HGSOC_06, HGSOC_08, HGSOC_14, and HGSOC_20) from patients with documented clinical backgrounds (Table S1). All PDO lines demonstrated stable growth patterns, maintaining stable stem cell potential *in vitro*, with expandability over 20 passages for more than six months (Fig. 1A).

**Figure 1.**
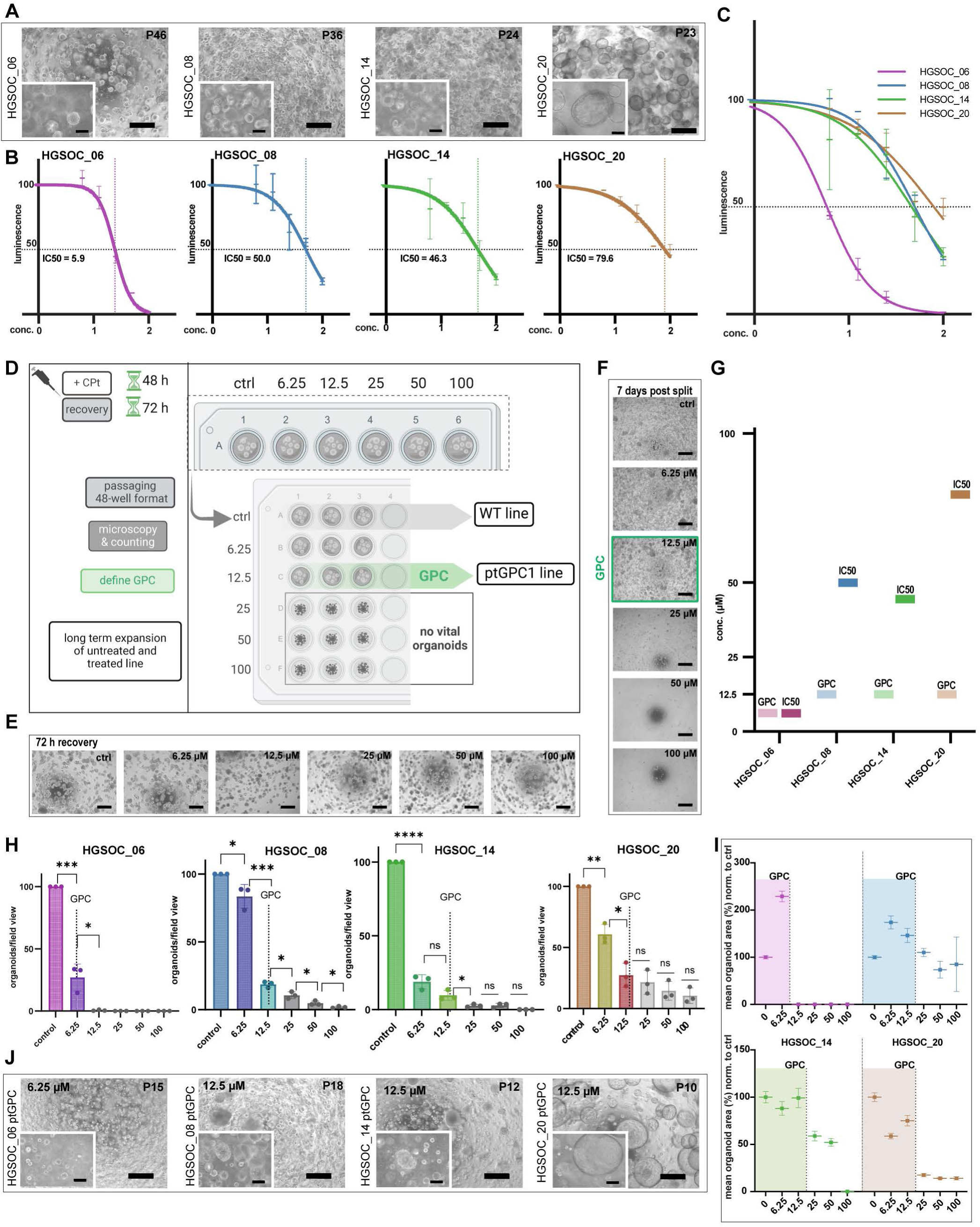
The Organoid Drug Resistance (ODR) test. (A), Phase contrast images of 4 PDO lines. Scale bars, 500 µm and 100 µm. (B), Individual and (C), Combined log-transformed dose-response curves showing patient-specific differences in sensitivity to carboplatin. Error bars are ± SEM of technical triplicates. Platinum IC50 concentration (µM). (D), Schematic ODR-test workflow illustrating key experimental steps of the assay. (E), Phase contrast images of representative PDO line at 72 h recovery and (F), 7 days post-split (P1) for each concentration (scale bar, 500 µm), green box = GPC for the given line, where long-term expansion was possible. (G), Direct comparison of GPC and IC50 levels in µM for each PDO line. (H), Diagram of normalized organoid counts in P1, Error bars are ±SEM of technical replicates from different wells. (I), mean organoid surface area diagrams showing the distribution of organoids that regrew after the carboplatin challenge. The range is calculated as mean ±SEM. (J), Phase contrast images of 4 ptGPC PDO lines, illustrating sustained long-term expansion >3 months. Scale bars, 500 µm and 100 µm.

Initially, we quantified the response of organoids to carboplatin by measuring the change in cell viability after 72 h, normalized to the control, and determined IC50 values for each patient-specific organoid line (Fig. 1B, C). The IC50 represents the concentration of carboplatin required to achieve a 50 % reduction of cell viability.

Notably, PDOs consist of cells at various stages of differentiation, with the progenitor population playing a key role in long-term cancer growth. Given that the residual growth potential of chemotherapy-surviving cancer cells presents the main challenge for strategies preventing recurrence, we focused on identifying sustained changes in organoids caused by sublethal carboplatin exposure. Therefore, we have designed the Organoid Drug Resistance (ODR)-test to identify and quantify the potential changes in stemness capacity as a consequence of drug treatment. In brief, PDO lines were exposed 48 h to carboplatin at five different concentrations (6.25 µM, 12.5 µM, 25 µM, 50 µM, and 100 µM) alongside a control, followed by a 72 h recovery phase and were subsequently passaged (P1) to a multiwell format in technical triplicates (Fig. 1D). The Growth Permitting Concentration (GPC) was defined as the highest concentration of carboplatin at which organoids maintained their stemness, assessed by their unlimited expansion potential, serving as a measure of patient-specific platinum resistance. While signs of acute cellular stress were notable right after the 72 h recovery (P0) (increase in granularity, dark appearance loss of adhesion, Fig. 1E and Fig.S1A), the functional impact of carboplatin on the organoid regeneration potential could be assessed only after singularization and reseeding (Fig. 1F). Compared to the previously determined IC50 values, the GPCs determined by the ODR-test were in three cases substantially lower (Fig. 1G). This implies that for these tumors a much lower platinum concentration is sufficient to exhaust long-term growth potential, than the one necessary to reduce overall viability to 50%. We observed a consistent trend of decreasing organoid formation potential with increasing concentrations of carboplatin in all donors. The number and size of the organoids in P1 were quantified by analysis of phase contrast images from independent wells in replicates. The magnitude of reduction in organoid count in P1 varied significantly between the lines, indicating substantial individual differences in their capacity to withstand platinum challenge (Fig. 1H).

The surface area of individual organoids was determined with the Qupath software annotation tool ^38^ (Fig. S1B, S1C). Surface size distribution analysis revealed that, despite a reduction in organoid formation efficiency, the mean size of the organoids that regrew in P1 remained stable until the GPC was reached (Fig. 1I). In line with this, at concentrations higher than the GPC, organoids were smaller and could not be further expanded (Fig. 1I, Fig. S 1D). Although initially viable, these 3D cellular aggregates effectively lost regeneration potential two to three weeks after carboplatin exposure. Importantly, organoids that regrew in P1 at GPC concentration despite a lower count in comparison to the control line could be successfully passaged in long-term culture (>3 months), with expansion potential comparable to that of *non-*treated controls (Fig. 1J). In conclusion, ODR testing determined a patient-specific critical concentration of platinum (GPC) required to effectively exhaust the stemness potential of HGSOC organoids. Average organoid size, rather than organoid count, after the challenge is a more suitable criterion to predict long-term growth capacity. Four ptGPC1 lines (post-treatment at GPC growing) from different donors were successfully established with continued stable growth in culture, confirming full preservation of growth capacity several months after carboplatin treatment.

### Phenotypic reprogramming in post-platinum organoids

The robust expandability of the ptGPC1 lines after the ODR-test provided the opportunity to compare them to respective untreated control PDO lines. The repair process of the double-stranded DNA breaks is coordinated by phosphorylated Histone protein A2X (yH2AX), initiating the recruitment of other repair proteins to the damage site, including BRCA1 ^39,40^. On average, more than two months passed since the exposure to carboplatin. Notably, ptGPC1 PDO lines still exhibited sustained elevated yH2Ax and pBRCA1 phosphorylation, as detected by confocal imaging, (Fig. 2A). Meanwhile, expression levels of stemness-related pathways, including Wnt and BMP target genes, as well as regulators of lineage fidelity and differentiation (PAX8, FOXM1, and CD133), analyzed by qPCR, showed no consistent changes in regulation among the different WT/ptGPC1 line pairs (Fig. 2B). The expression levels of ID2 and ID3, which are downstream of active BMP signaling, remained relatively stable and robust, as indicated by low ΔCt values (Fig. S 1E). This aligns with the preserved growth capacity of ovarian cancer PDOs, as BMP activity is known to promote their growth ^8,12^. Despite preserved growth capacity, ptGPC1 organoids showed changes in cellular architecture visualized by actin stain phalloidin as well as prominent defects in the morphology of the nuclei (Fig 2C). The expression pattern of the nucleolus marker fibrillarin was found to be a more condensed and larger area in comparison to controls (Fig. 2D). These findings suggest that carboplatin induces irreversible molecular changes in organoids without impairing the general ability of cell proliferation and progenitor potential.

**Figure 2.**
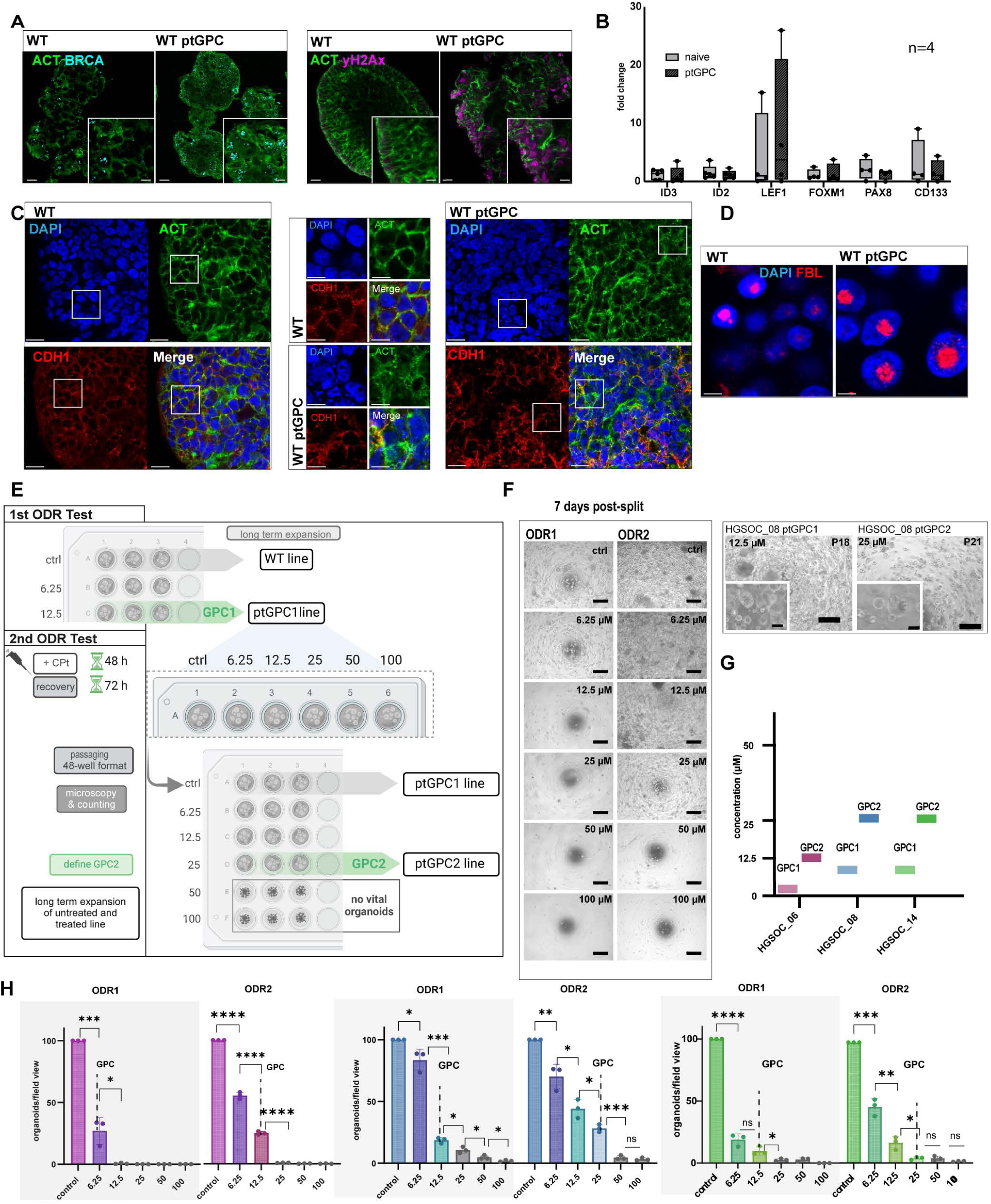
*In vitro* modeling of platinum resistance. (A), IF staining of p-BRCA (left) and γ-H2Ax (right) of untreated and ptGPC PDOs showing sustained DNA damage and activation of DNA repair in pre-exposed lines. Scale bars, 20 µm and 10 µm. (B), qPCR of expression of stemness- and differentiation-related target genes in ptGPC organoids (n=4 independent biological sample pairs). Error bars ± SEM. (C),(D), IF staining showing changes in the organization of cytoskeleton (phalloidin, green), nuclear morphology (DAPI, blue), and nucleolus (fibrillarin, red) in untreated vs. ptGPC PDOs. Scalebar, 20 µm, 10 µm and 5 µm. (E), Schematic ODR2-test workflow for the rechallenge of the ptGPC1 organoids. (F), Direct comparison of 7 days P1 phase contrast images for the representative donor line after ODR1 and ODR2 test, showing differences in sensitivity/growth potential (scale bars, 500 µm, and 100 µm) leading to (G), increasing GPC (GPC1 to GPC2 comparison in µM), for three pairs of PDO lines. (H), Quantification of the change in drug sensitivity, measured by organoid count, normalized to the control for ODR1, and ODR2. Scale bars are ±SEM of technical triplicates, and P-values are calculated by the Student t-test (*p <0,05, **p<0.01; ***p<0.001, **** p<0,001).

To further evaluate whether the observed cellular phenotype in ptGPC1 organoids plays a functional role in response to the platinum challenge, we subjected them to a second round of carboplatin treatment and ODR testing (ODR2-test) using the same workflow (Fig. 2E). We found that organoids that underwent re-challenge showed notably improved formation and growth potential after the second treatment when compared with the outcomes from ODR1-test (Fig. 2F, Fig. S 2A-D). For all tested lines the growth-permitting concentration of the second challenge - defined as GPC2 - surpassed the previous GPC1 value (Fig. 2G). Importantly, for each line and at every carboplatin concentration, organoid counts were consistently higher post-ODR2 compared to post-ODR1 counts resulting in enhanced ODR-response profiles (Fig. 2H). In summary, repeated *in vitro* platinum treatment and ODR testing revealed the gradual development of resistance in PDO organoids, including those derived from patients with clinically favorable platinum responses.

### Cytoskeleton remodeling promotes platinum resistance

Next, we aimed to gain a deeper understanding of the biology of ptGPC organoid lines by evaluating their transcriptomic and proteomic landscapes, to obtain a global view of the therapy-driven long-term changes. Bulk RNA sequencing of two paired donor PDO lines (HGSOC_06 and HGSOC_08), with additional biological replicates (*non*-treated and ptGPC lines) revealed a strong patient-specific component in the expression profiles, as visualized by principal component analysis (PCA), with carboplatin treatment not having a defining impact on the overall gene expression profile (Fig. 3A). This great interpatient heterogeneity is in line with previously published scRNA data from ovarian cancer tumor tissue ^41^. Differential expression analysis performed with the DESeq2 R package identified 837 genes differentially regulated in ptGPC1 lines (p-value <0.05), among which 80 were significant in adjust < 0.05 analysis (Table S2). The Volcano plot of the differentially expressed genes (*non*-treated vs. ptGPC lines) revealed the most strongly regulated candidates (Log2Foldchange, and p-value) in response to chemotherapy (Fig. 3B). Despite a pronounced difference between the two donors, reflected also in their clinical background, the set of genes showed a consistent pattern of change in expression driven by carboplatin treatment. As illustrated by heat-map analysis of the top 25 regulated genes based on their significance (p-adjust < 0.05), samples clustered according to the treatment, confirming the existence of a common signature response of the organoids upon carboplatin exposure (Fig. 3C). The Gene Set Enrichment Analyses (GSEA) of regulated genes for GO: Biological Processes revealed significant positive enrichment or upregulation in gene sets involved in tissue and epithelial development and differentiation, while cell adhesion regulators were found downregulated in ptGPC1 organoids (Fig. 3D). Mass spectrometry analysis of the protein lysates of the same pair of organoid lines in quadruplicates confirmed key findings of the transcriptional profiling, with highly similar sample distribution in the PCA plot (Fig. 3E). Likewise, GSEA for Biological Processes showed significant enrichment in pathways regulating migration and cell adhesion (Fig 3F). On the proteome level, a total of 947 (p-adjust < 0.05) proteins were confirmed to be regulated in the same fashion in both donors (Table S3). The enrichment score plots (ES) for the downregulation of adhesion-related signatures in transcriptomic and proteomic data sets yielded comparable results providing complementary evidence obtained by independent omics methodologies of the downregulation of adhesion complexes in ptGPC organoids (Fig 3F, G).

**Figure 3.**
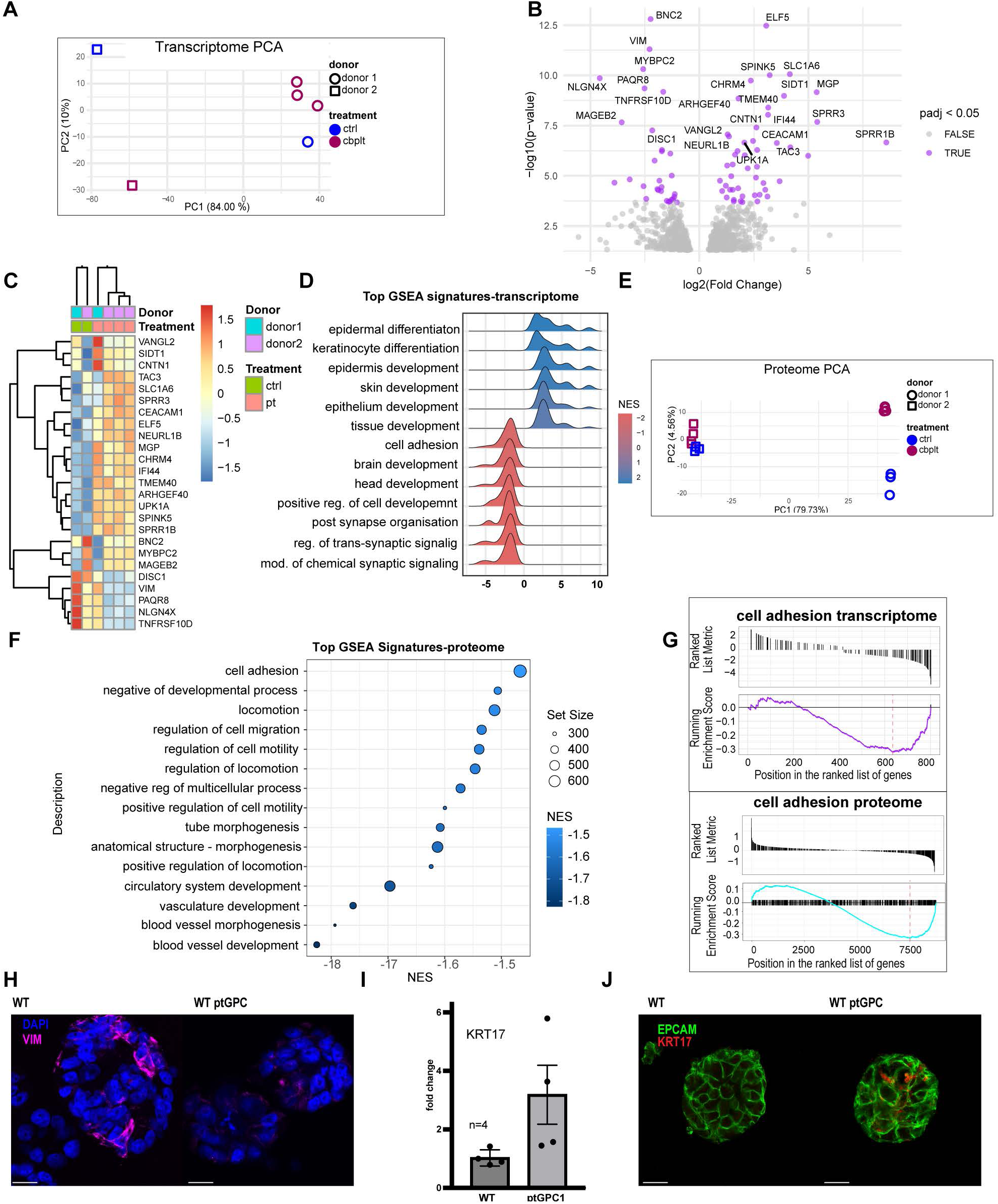
Post-Platinum organoid transcriptomic and proteomic characterization. (A), PCA plot of mRNA sequencing of control vs. GPC lines, for two donors, showing patient-specific characteristics dominating expression profiles of the organoids. (B), Volcano plot of top 25 regulated candidate genes (Log2 Foldchange, p-adjust < 0.05). (C), heat-map analysis of the top 25 regulated genes (p-adjust < 0.05), illustrates that despite great differences between donors samples cluster according to treatment. (D), GSEA for GO Biological processes in untreated vs. ptGPC1 lines identifies enrichment in pathways regulating cytoskeleton and differentiation. (P-adjust< 0,05). (E), PCA plot of mass spectrometry measurements (n=4 independent technical replicates for two sets of lines ctrl vs GPC, total 16 samples), confirms that patient-specific characteristics are defining components in the global proteome profile. (F), GSEA of top enriched pathways for proteome (NES), identifies enrichment in downregulation for pathways regulating cell adhesion and locomotion. (G), ES plots for cell adhesion in transcriptome and proteome data. (H), Loss of vimentin IF staining visualized by confocal imaging. ptGPC1 organoids in comparison to the control line. Scale bar, 20 µm. (I), qPCR validation of the induction of KRT17 in ptGPC1 organoids (n=3 independent biological samples) and (J), by IF staining in confocal imaging. Scalebar, 20 µm.

Remodeling of the cytoskeleton and changes in adhesion have previously been found to promote therapy resistance. Downregulation of vimentin, one of the main regulators of mechanosensing ^42^ has been reported in ovarian cancer cell line models during the acquisition of resistance ^43,44^. Accordingly, a reduction of vimentin signal intensity was detected by confocal imaging in our organoid ptGPC model (Fig 3H), validating the RNA sequencing data, since vimentin was one of the top downregulated genes as visualized in the heat-map (Fig. 3C).

Considering the significant changes in genes regulating cytoskeletal organization detected in the transcriptome of ptGPC lines, along with previous studies linking KRT17 to the stemness compartment of the fallopian tube and ovarian cancer ^17–20^, we investigated whether KRT17 has a functional role in the development of platinum resistance. Though initially below the (p-adjust) threshold in bulk RNA profiling, transcriptional upregulation in KRT17 organoids in response to carboplatin and subsequent ODR-testing was confirmed by qPCR for four independent ptGPC1 paired samples of different donors (Fig. 3I). Congruently, a rise in KRT17 levels in ptGPC1 organoids were visualized by confocal imaging (Fig. 3J).

### Functional KRT17 overexpression increases the platinum resistance of the organoids

To assess the functional relevance of KRT17 in the process of increasing resistance, we edited the KRT17 gene locus using CRISPR/Cas9, introducing specific gRNAs by nucleofection, followed by the generation of a monoclonal line. Editing resulted in the efficient excision of 96 bases in exon 4, as confirmed by Sanger sequencing (Fig. S 3A-C) qPCR analysis of the expression of individual exons of the KRT17 gene, in the crKRT17 HGSOC_08 line confirmed the complete loss of exon 4 expression, while all other tested exons showed preserved transcription levels (Fig. 4A). In alignment, Western Blot revealed a truncated KRT17 protein at ∼ 40 kDa, with the absence of a full-length protein in the CRISPR/Cas9-engineered donor line (Fig. 4B).

**Figure 4.**
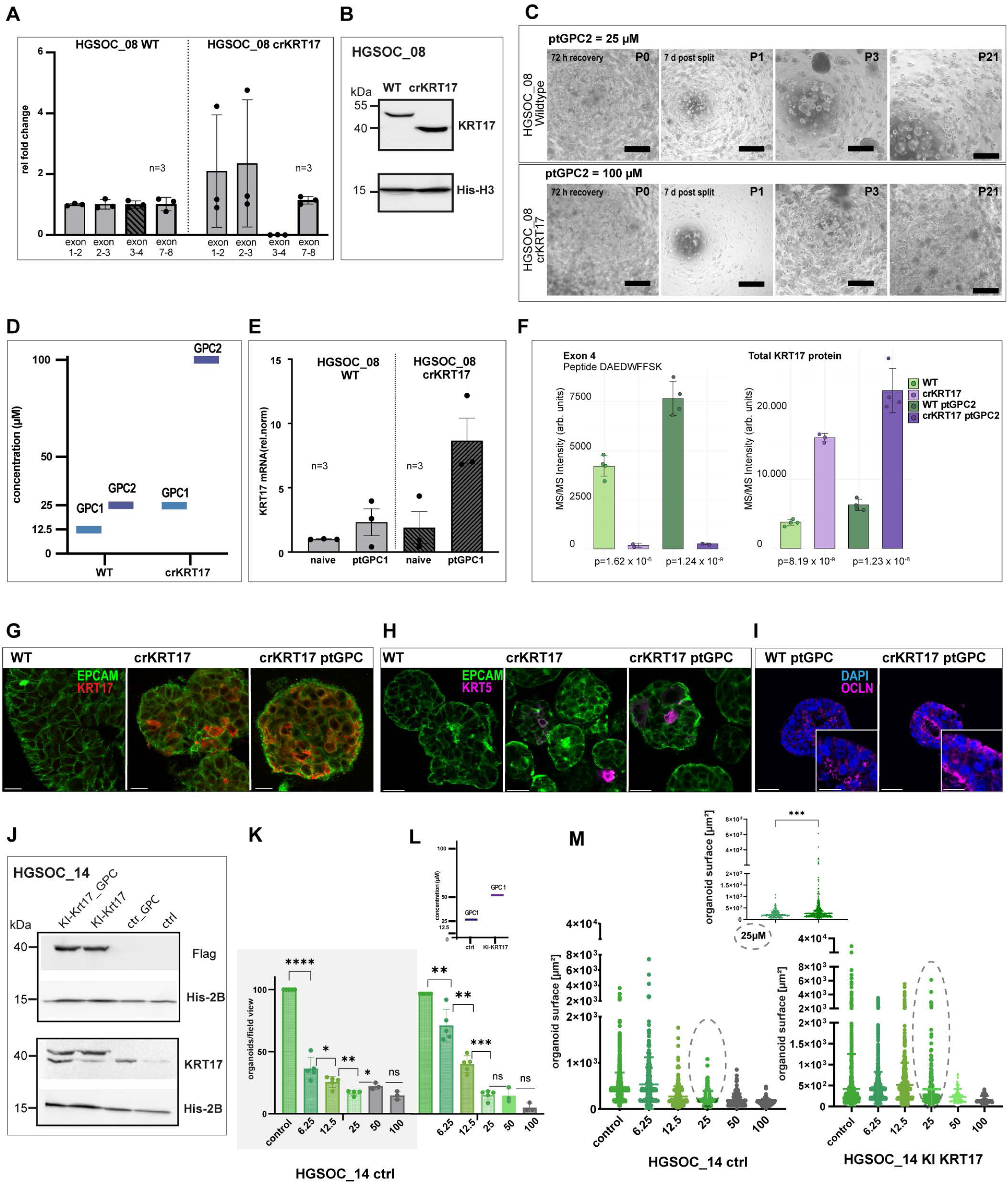
Platinum hyper resistance by KRT17 editing. (A), qPCR of individual KRT17 exons, in control vs. crKRT17 edited lines shows loss of expression of exon 4 (n=3 independent biological replicates). Errorbar ±SEM. (B), Western blot for KRT17 confirms expression of the truncated protein. (C), Comparative phase contrast images after ODR2-test for WT crKRT17 lines illustrating long-term expansion potential after sustaining 25 µM and 100 µM challenge in P0. Scale bars, 500 µm. (D), Bar plot summarizing shift in GPC levels and a strong increase in crKRT17 line. (E), KRT17 RNA expression by qPCR of treated and untreated, WT and crKRT17 lines, illustrates strong induction in the expression of KRT17 in crKRT17 organoids (n=3 independent biological samples). Errorbars ±SEM. (F), Mass spectrometry data show a significantly lower or absent MS/MS intensity of the DAEDWFFSK peptide, corresponding to the coding region of exon 4, in edited organoid lines compared to total KRT17 protein intensity (n=4 independent biological samples), confirming the overexpression of the remaining portion of the protein in ptGPC2 organoids. (G), Confocal imaging validates a strong increase in KRT17 levels and (H), KRT5 levels. Scalebars, 20 and 50 µm. (I), Re-distribution of Occludin staining signal to lateral membranes in crKRT17 ptGPC2 organoids indicates loss of polarity. Scalebar, 50 µm and 25 µm. (J), Western blot for KRT17 expression in KI-KRT17 and control lines, with Flag and endogenous KRT17 antibody, confirms expression of the exogenous construct as well as induction of endogenous protein in response to carboplatin. (K), Phase contrast images of control and KI-KRT lines at two different carboplatin concentrations (25 µm and 50 µm) at P1. Scalebars, 100 µm. (L), Quantification of the ODR1-test of KI-KRT17 vs. lenti-ctrl organoids showing the increase in organoid counts and rise in GPC. GPC1=25 µM (lenti-ctrl) and GPC1=50 µM (KI-KRT17). Scalebars, 100 µm. (M), Analysis of the distribution of organoid sizes reveals a group of significantly larger organoids (dotted circles) in ptGPC KI KRT17 culture suggestive of increased stemness potential. Error bars ±SEM of measurements in independent wells. P-values are calculated by the Student t-test for the quantification of organoid counts, and the Mann-Whitney U test for comparison of differences in organoid sizes (*p <0,05, **p<0.01; ***p<0.001, **** p<0,001).

We then investigated whether truncation of the KRT17 protein influences the response of organoids to carboplatin by ODR testing. Compared to the WT ptGPC lines, crKRT17 ptGPC lines exhibited strongly increased resistance to consecutive carboplatin treatment (two rounds of ODR-testing), as organoids regrew in P1 following the exposure to maximal (100 µM) carboplatin concentration (Fig. 4C, Fig. S 3D, E). Consequently, GPC1 and GPC2 levels were higher in the crKRT17 line (GPC1 25 µM, GPC2 100 µM), when compared to the WT line (GPC1 12.5 µM, GPC2 25 µM) (Fig. 4D). Notably, quantification of the KRT17 gene expression level in crKRT17 ptGPC2 organoids compared to WT controls detected a strong upregulation in the transcripts from remaining exons following repeated carboplatin treatment, suggesting the existence of synergistic mechanisms between the effect of carboplatin and regulation of the KRT17 gene locus (Fig. 4E). To confirm the upregulation of KRT17 at the protein level and specific truncation at the peptide level, we performed a DIA-based proteomics experiment. The results showed an overall increase in total KRT17 intensity in crKRT17 *non*-treated and ptGPC2 lines in comparison to WT counterparts which fully aligned with qPCR data. Notably, we found one peptide DAEDWFFSK, that corresponds to the genomic deletion in exon 4, that has significantly lower or absent MS/MS intensity in the edited line (Fig. 4F) (p-value =1.6 ×10^-6^ and p value=1.24×10^-9^ for control and ptGPC lines respectively). At the same time, platinum treatment further strongly boosted levels of all other KRT17 peptides in the crKRT17 lines (total protein) (p-value=8.19×10^-9^ and p=1.23×10^-8^), and upregulation was confirmed for all individual exons (Fig. S 3F). An additional strong increase in KRT17 levels, triggered by edit was also confirmed by confocal imaging (Fig. 4G). Further, hyper-resistant crKRT17 ptGPC2 organoids exhibited an increase in KRT5 expression, a mediator of dedifferentiation and marker of epithelial stages of increased plasticity ^45^, as well as in the distribution of the occludin staining to basolateral membranes indicating further loss of polarity (Fig. 4H, I).

As CRISPR/Cas9 editing failed to achieve full knockout of KRT17 suggesting the potential existence of selection pressure favoring edits where gene expression is partially maintained, we have chosen a complementary approach and alternative strategy to prove the direct ability of KRT17 to modulate carboplatin response. Full-length (Myc-DDK tagged) KRT17 was introduced by lentiviral transduction (knock-in construct, KI) into a different PDO line (HGSOC_14) to additionally exclude donor-specific effects (Fig. S 4A.) The protein expression from the KRT17 open reading frame was validated by both Flag-antibody detecting the tagged protein (Fig. 4J) as well as with KRT17 antibody detecting both endogenous and slightly larger Flag-Myc tagged KRT17 protein (double band). ODR-testing revealed that expression of the exogenous KRT17 leads to a strong increase in the organoid count, in comparison to the organoids transduced with the control vector (Fig. 4L). In agreement with this, GPC1 shifted from 25 µM to 50 µM in KRT17 (Fig 4L) transduced organoids confirming the ability of KRT17 to directly increase platinum resistance. Interestingly carboplatin treatment in the range of 12.5-25 µM, led to the appearance of significantly bigger, larger organoids in Flag KRT17 ptGPC1 lines in P1 (Fig. 4M), which subsequently translated into a strong increase in the growth rate of the KI-KRT17 organoids during subsequent passaging, quantified by splitting ratio (Fig. S 4C)

### K-Score is an independent negative prognostic factor of OS and PFS in HGSOC patients

To investigate the association of KRT17 expression with clinical outcomes, we have conducted an exploration analysis of clinical patient data to quantify the expression frequency of KRT17 across a high number of HGSOC patients and to evaluate its prognostic impact. Therefore, tissue-microarrays (TMA) of 384 patients diagnosed with primary HGOSC at advanced disease stages (FIGO III-IV) were stained for KRT17 by immunohistochemistry. Clinical characteristics and descriptive statistics of the patient cohort are listed in Table S4 and Table S5.

The evaluation of the KRT17 expression was conducted by digital AI-based imaging analysis ^38^ (Fig. S 4D). To quantify expression levels, we developed the K-score as a measure of the KRT17 expression, which is based on the number of positively stained tumor cells in proportion to the overall detected tumor cells and ranged between 0-200 % (Fig. 5A, Table S6). The average KRT17 expression across all tested tumors was generally low, with a mean K-score of 2.82 % (Fig. S 4E). By using the biomarker cutoff finder tool ^46^, we defined a K-score of 5 % as a benchmark to separate the populations of KRT17^high^ and KRT17^low^ tumors. In Kaplan-Meier-analysis, the median overall survival for KRT17^high^ patients was 21 months (95% CI 35.3-46.7 months) compared to 41 months (95% CI 11.1-30.9 months) for KRT17^low^ patients (p=0.0043) (Fig. 5B). Median progression-free survival for KRT17^high^ patients was 14 months (95% CI 10.9-17.2 months) compared to 19 months (95% CI 17.1-20.9 months) for KRT17^low^ patients (p=0.0081) (Fig. 5C). Cox regression multivariate analysis, controlled for the main factors influencing the prognosis (general health status of the patients (ASA), age at first diagnosis, postoperative residual disease and neoadjuvant chemotherapy) showed that the K-score is an independent prognostic factor on overall survival with a Hazard ratio of 1.84 (CI 1.17-2.87) (Fig. 5D) and progression free survival with a Hazard ratio of 1.58 (CI 1.06-2.34) (Fig. 5E). Taken together, the data consistently showed that KRT17 not only has a functional role in the cellular response to platinum in organoids, but its expression level in tumor tissue has prognostic significance in a large cohort of advanced HGSOC patients, thus, making it a possible target for novel therapeutic approaches.

**Figure 5.**
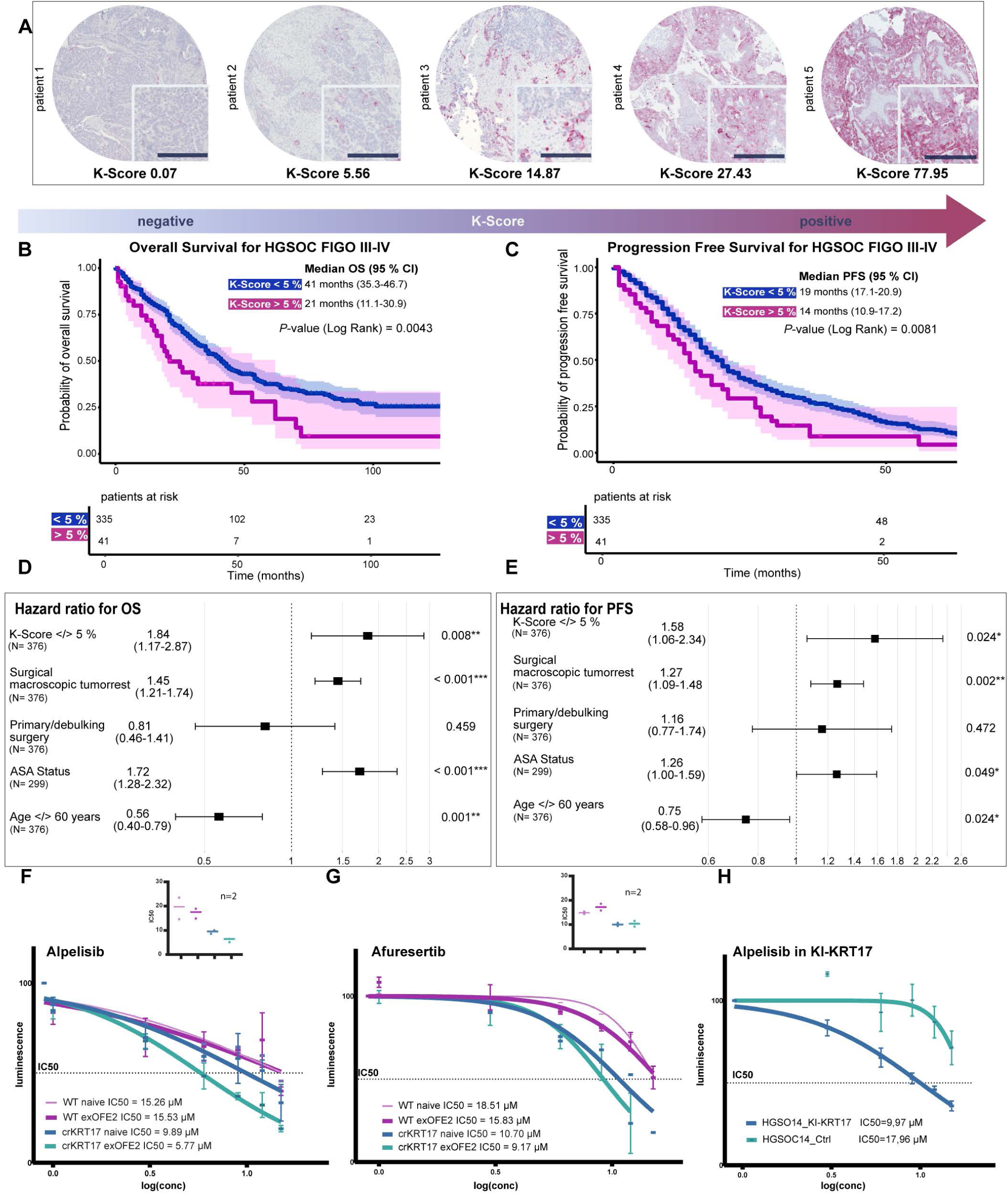
Prognostic K-score as stratification to PIK3/Akt-targeted therapy. (A), Images of KRT17 immunohistochemical staining representing TMA cores of 5 exemplary patients displaying different levels of expression. (B), Kaplan-Meier analysis in the patient cohort of advanced HGSOC cases. K-score cutoff, 5 %. Reveals significantly shorter OS (p-value (Log-Rank) = 0.0043) and (C), PFS (p-value (Log-Rank) = 0.008) for the group of KRT17^high^ patients. (D), Multivariate Cox regression analysis for OS and (E), for PFS, confirms the prognostic value of the K-score as an independent risk factor (HR 1.84, p-value=0.008 for OS, and HR 1.58, p-value=0.024) for PFS. (F), log-transformed dose-response curves for alpelisib in untreated and carboplatin-treated WT and crKRT17 lines, show a strong increase in sensitivity in the lines expressing elevated KRT17 levels. (G), IC50 values in µM, in independent biological replicates of the assay and (H), (I), showing results of the analogous testing setup for afuresertib. (K), Alpelisib treatment of the KI-KRT17 expressing independent donor line confirms the capacity of KRT17 to directly modify the sensitivity of the organoids to PI3 kinase pathway inhibition. Error bars are ±SEM of technical triplicates.

### KRT17 is a potential stratification marker for PI3K/Akt-pathway-targeted therapies

KRT17’s involvement in cellular stress response by activation of downstream signaling, including the PI3K/Akt pathway has previously been reported in ovarian cancer models ^23,37^. By now, the PI3K-inhibitor Alpelisib and pan-Akt-inhibitor Afuresertib have shown tolerable safety profiles for the treatment of HGSOC at the recurrent platinum-resistant stage but failed to reach primary endpoints in studies designed to include all-comer study populations ^47–49^. We tested whether a high KRT17 expression might be a potential stratification factor to determine which patients might benefit from PIK3-Kinase- and/or Akt-inhibition. Therefore, we have exposed to platinum WT non-treated control PDOs and WT ptGPC2 PDOs, crKRT17 platinum *non*-treated control PDOs, and crKRT17 ptGPC2 PDOs with the PI3-kinase- and Akt-inhibitors Alpelisib and Afuresertib.

CrKRT17 *non*-treated control PDOs, overexpressing KRT17, were found to be more sensitive to Alpelisib (IC50 9.89 µM vs. controls IC50 15.26), and the crKRT17 ptGPC2 PDOs showed the lowest IC50 value was stable and reproducible in independent set of experiments (IC50 plot) (Fig. 5F). In the case of Afuresertib, high levels of KRT17 induced sensitivity already in the crKRT17 *non*-treated control PDO line (IC50 10.7 µM), without further reduction in the crKRT17 ptGPC2 line (IC50 9.17 µM, Fig. 5G). Finally, we have validated the role of KRT17 in the response of organoids to Alpelisib, by treatment of the KI-KRT17 line in comparison to control-transduced organoids which confirmed a strong increase in sensitivity to Alpelisib (Fig. 5H).

These results demonstrated that elevated KRT17 levels and previous exposure to chemotherapy predispose tumors to effective inhibition of PI3K signaling by Alpelisib, while pan-Akt inhibition by Afuresertib primarily targets the KRT17 status irrespective of the carboplatin effect. In conclusion, the KRT17 expression defined by the K-score status could be a valuable stratification tool to select patients who could benefit from Akt-inhibitor treatment in advanced and platinum-treated HGSOC.

## Discussion

In the present study, we developed the ODR-test as a novel and innovative experimental approach to quantify platinum resistance in ovarian cancer and study cellular plasticity which is driven by the response to therapy. Although different groups previously reported an association between drug resistance and enrichment of stemness-related phenotypes within tumors, the mechanism of how chemotherapy resistance emerges remains poorly understood ^15,50,51^. Since organoid growth in culture is based on the preservation of epithelial progenitor potential, the ODR-test determines and quantifies the conditions under which individual tumors can sustain growth capacity despite the damaging effects of carboplatin. This represents an important conceptual advancement to commonly used drug assays as they only determine the acute and direct toxic effects of the drugs on remaining viable cells ^13,52^. Clinical experience has long suggested that every ovarian cancer patient develops molecular resistance to repeated platinum treatment over time, even though the extent of resistance and the speed of the process may vary between patients and could not be translated into a clinically applicable biomarker stratification so far ^5^. Importantly, the ODR-test effectively confirmed the existence of this phenomenon *in vitro*, as organoids consistently exhibited an increase in resistance upon re-challenge measured by a shift in the organoid count and an increase in growth permitting concentration (GPC) regardless of their clinical outcome. The data is also fully consistent with the hypothesis that carboplatin treatment triggers a conserved mechanism of the cellular response while the dynamics of the resistance acquisition have a strong individual component that can be analyzed in detail in patient-specific lines. Different degrees of resistance, observed in organoids after repeated ODR testing also explain why increase does not carry clinical consequences in all cases, as the basal level of sensitivity of each tumor is likely the result of complex genetic and environmental factors, and good responders remain more sensitive despite a molecular drift towards a resistant phenotype.

Consequently, the ODR-test could be a useful tool to successfully address the challenge of identifying patients at high risk for resistance development ^4^. Global analysis of gene expression and the proteome of ptGPC lines revealed changes in expression in cellular networks regulating cell adhesion, cell-cell communication as well as cytoskeletal components (Figure 3A-G). The similarities in response of different donor lines of variable basal resistance to carboplatin challenge are evidence of conserved reprogramming mechanism and cellular plasticity under therapeutic pressure. While more research is needed to fully understand the mechanisms of stemness preservation under platinum treatment, the stable expansion of ptGPC lines offers a new valuable model to study this phenomenon. Identification showing that KRT17 is responsive to cytotoxic stressors such as platinum-induced DNA damage ^53^. The strong increase in platinum resistance profile that we detected in lines expressing elevated levels of KRT17 by either CRISPR/Cas9 editing of exon 4 or through genomic integration of additional copies of the KRT17 gene, demonstrated the ability of KRT17 to directly influence the response to carboplatin and represents evidence of a potential regulatory loop. Altogether, these findings support the hypothesis, that platinum-caused DNA damage induces KRT17, which subsequently actively participates in changes in the cytoskeletal organization to enhance cellular stemness, promoting tumor survival ^23,34,37^. The clinical prognostic relevance of KRT17 expression was established by immunohistochemical analysis of the KRT17 expression profile in a large cohort of 384 advanced HGSOC patients. We found that the KRT17 expression level classified by the immunoreactive K-score, has a significant prognostic value for both overall and progression-free survival. This clear prognostic relevance, confirmed by the Cox proportional hazards model in combination with the *in vitro* results on organoids, supports further studies and the development of the K-score as a biomarker in a clinical context to determine which patients are likely to develop a platinum-resistant recurrent disease. As a high KRT17 expression has been reported to have negative prognostic significance also in other epithelial cancer entities, our study could be of interest in a broader oncological context ^24–34^.

Genetically modified organoids expressing high levels of KRT17 (ptGPC crKRT17 and KI KRT17 lines) showing high levels of platinum resistance were a suitable model to study the response of ovarian cancer to targeted agents as they reflect the potentially relevant clinical scenario of heavily pre-treated, platinum-resistant, and highly KRT17-expressing tumors. In these organoids (Fig. 5F, G, H), we have identified a strong improvement in sensitivity to the pan-Akt-inhibitor Afuresertib and PI3K-inhibitor Alpelisib. This direct relationship between KRT17 expression and sensitivity of organoids to Akt/PI3 kinase inhibition strongly suggests that further development of KRT17 as a promising biomarker and stratification tool to implement therapies targeting Akt/PI3-kinase in the treatment of platinum-resistant, recurrent ovarian cancer is warranted.

Patient-derived organoids effectively capture in vitro basic organization of the epithelial compartment of the tumor tissue, as they expand in culture based on the balance of stemness and differentiation mechanisms governing tumor growth. Though principal findings of the ODR testing and analysis of ptGPC lines suggest that the model has sufficient complexity to phenocopy gradual rise in platinum resistance observed in patients, the absence of immunosurveillance and tumor microenvironment likely limits the predictive potential of the model regarding the timing of the disease recurrence, and exact timing when the tumor will reach a resistance level which would be consequential for clinical decision making. In line with this, while the K-score is significantly associated with negative patient outcomes, it represents an indicator of a faster process of developing resistance during disease progression.

## Acknowledgments

Research at the Research Laboratory of the Department of Obstetrics and Gynecology is made possible by the institutional support of the LMU University Hospital. M.K., S.M., F.T., A.C.R, B.C obtained the support of the German Cancer Consortium (DKTK). J.R and A.C.R received funding from Bavarian Cancer Research Center BKZF. J.R and A.C.R obtained fellowships of the LMU Medical and Clinician Scientist Programm. F.T. received grant funding from German Cancer Aid DKH, # 70113426 and #70113433.M.K, J.O, I.P, and S.G had the support of the ERAPERMED2022-141 (BMBF Grant 01KU2302), I.P and J.V.O work at the Novo Nordisk Foundation Center for Protein Research (CPR), The authors would like to thank Linpu Yang, Maria Fischer, Andrea Sendelhofert, Martina Rahmeh, Madlen Leinauer, Cornelia Herbst, Sabine Fink, and Simone Hofmann for their help. We are grateful to the staff of BMC Core Facility for Bioimaging, Klein Lab at Dr. von Hauner Childreńs Hospital, who provided S2 Lab space, and the Core facility of the Institute of Anatomy, LMU Faculty of Medicine, where organoid samples were embedded.

## Author Contributions

Conceptualization, M.K, and J.R.; methodology J.R, M.K., I.P, J.V.O; validation J.R., J.S, S.H, T.Z, S.G.; formal analysis J.R, J.S, I.P., S.H., S.G, T.Z, A.D., J.V.O and M.K.; investigation J.R., J.S., S.H., T.Z., I.P, S.G, J.F., A.C.R., A.D., and N.V.; resources S.M., C.K., F.T., A.B, B.C., F.K., D.M, T.S.; writing-original draft J.R., M.K.; writing-reviewer and editing J.R, M.K., F.T, S.M., F.K., B.C., D.M., A.B., T.S., C.K., I.P., J.V.O., S.G., J.S.; visualization J.R, J.S, S.G, I.P, and M.K.; supervision M.K. project administration M.K.; funding acquisition M.K., F.T., J.R., A.C.R., J.V.O.

## Methods

### Experimental model and study participants

#### Human subjects

The study was conducted following the Ethics Committee of the Ludwig-Maximilian-University (LMU), Munich, Germany (17-0471), and in compliance with all legal EU, national, and local regulations. Each patient provided written informed consent to participate. Ovarian cancer biobank, established following our institutional protocols ^12,54^ was a source of primary high-grade serous ovarian cancer patient-derived organoid (HGSOC PDO) lines that were used in this study including embedded tissue samples, along with their clinical data. A routine pathological assessment of the primary tumor tissue confirmed the malignant, non-borderline diagnosis based on standard histomorphology characteristics. The samples were collected during multi-visceral surgeries at the Department for Gynecology and Obstetrics, LMU Hospital, Munich, Germany. Our gyneco-oncological surgeons collected clean peritoneal tumor deposits, avoiding macroscopically necrotic areas, and the tissue was processed in our laboratory for organoid generation, as previously described ^12,54^. Embedded tissue and corresponding organoid cultures exhibit matching molecular profiles ^8^.

Tissue microarrays (TMA) for immunohistochemical analysis were prepared and stained by the LMU Department of Pathology, comprising a total of 384 duplicated samples of HGSOC patients diagnosed with advanced staged disease (FIGO III-IV) treated at our department. Clinical data was available for further analysis.

#### Organoid cultivation

HGSOC PDOs were cultivated in individually optimized media according to the previously described protocol. Donor lines included in the study all grew in OCM1 medium (low WNT, high BMP environment) with two lines showing benefit from the addition of her 1β (Table S7) ^12^. Passaging of organoids cryopreservation and thawing procedures followed protocols established by our laboratory^12^. Briefly, depending on organoid growth, every 10-20 days, organoids were released from Cultrex® BME 2 and trypsinized with TrypLE™ (Gibco^TM^). Depending on the grade of the desired dissociation, the cell suspension was strained through cell filters to ensure singularization. Dissociated cells were re-seeded in matrix ovarian cancer medium was added after 30 min of solidification. Cryopreservation of organoids was performed in the proliferation phase of organoid growth in the first week *post*-passage Organoids were pelleted and transferred to 1 ml Cryopreservation medium (e.g., Freezing Medium Cryo-SFM, PromoCell – Human Centered Science®) and stored initially at −80°C freezing container (e.g. Nalgene® Mr. Frosty, Sigma-Aldrich®), before transferring to liquid nitrogen for long-term stable storage.

#### *In-vitro* Cell viability drug assay

Organoids were trypsinized, strained through the filter, and counted and 20-30,000 cells were seeded in in a 48-well plate format in triplicates. At day 5-7 *post*-seeding, depending on the growth dynamic of the individual line, fully formed organoids were treated with carboplatin at increasing concentrations (dilution in organoid medium to 6.25 μM, 12.5 μM, 25 μM, 50 μM, and 100 μM), alongside a control. The cell viability was measured after 72 h using the CellTiter-Glo® 3D Cell Viability Assay (Promega) according to the manufactureŕs directions at the Varioskan™ LUX multimode microplate reader (Thermo Scientific™). Drug response for PI3K/Akt inhibitors Alpelisib and Afuresertib was measured in the same format with a treatment duration of five days. The individual inhibitory concentration at 50 % cell viability was determined for each tested PDO line, in Graph Prism software by calculating dose response by applying a nonlinear regression model.

#### Organoid Drug Resistance (ODR) test

The regenerative potential of organoids was tested by subjecting them to different doses of carboplatin using our developed ODR test (Fig. 1D). Initially, organoids were singularized by trypsinization, mechanical dissociation, and straining through a 200 µm filter for equal distribution and seeded at density 50,000 cell/well in a 24-well plate format. At day 7 *post*-passage, the organoids were treated with carboplatin for 48 h at five increasing concentrations (dilution in organoid medium to 6.25 μM, 12.5 μM, 25 μM, 50 μM, and 100 μM, alongside a control. After a 72-hour recovery phase in a carboplatin-free organoid medium, passaging was performed to a 48-well format in a 1:3 ratio into Passage 1 (P1). Organoid generation and growth were quantified by imaging each well at 4x and 10x magnifications using a phase contrast microscope, at 3-, 7-, and 10-days post-split, and using the annotation tool of Qupath Software. By further passaging of organoids, we identified the highest carboplatin concentration level in P0 after which unlimited long-term expansion of the line was possible permitting concentration (GPC).

#### RNA Isolation

The RNeasy Mini Kit (QIAGEN) was used for total RNA extraction according to the manufactureŕs protocol. Organoids were released from the extracellular matrix with cold ADF++ and pelleted. After resuspension in Buffer RLT (QIAGEN) supplemented with ß-mercaptoethanol and homogenization 70%-ethanol was added, and samples were processed via RNA purification columns. Quantification of the RNA concentration and purity was performed with the Qubit 4 Fluorometer (Thermo Scientific™) following the instructions of the Qubit™ RNA HS Assay Kit (Thermo Scientific™).

#### Reverse Transcription and qPCR

For the two-step quantitative reverse transcription PCR (qRT-PCR), the RNA samples were transcribed to (cDNA) using the cDNA Synthesis Kit (Biozym). TaqMan® Universal PCR Master Mix, TaqMan® Gene Expression Assay targeting the gene of interest (Table S8) was used for measuring quantitative gene expression on the Applied Biosystems™ 7500 Fast Real-Time PCR System. The quantification of expression levels was performed by ΔΔ Ct method relative to the expression of the Glycerinaldehyd-3-phosphat-Dehydrogenase (GAPDH). Change of expression in the test group (e.g. ptGPC organoids) was determined in relation to the average value of the expression of the gene in the control group. The GraphPad Prism 10 software was used for data analysis and graphical representation.

#### CRISPR-Cas9 mediated editing of organoids

CRISPR-Cas9 mediated gene editing for KRT17 was performed with two customized single guide RNAs (sgRNA) (Synthego) (Table S9). Organoids previously singularized by trypsinization and mechanic dissociation were transfected with the Cas9 endonuclease and sgRNA /ribonucleoparticles (RNP) via electroporation in the NEPA21 electroporator (Nepagene). For this purpose, 150,000 singularized organoid cells each were combined in a total of 50 µl reaction mixture consisting of 25 µL reduced serum medium (Opti-MEM^TM^, Gibco^TM^) and 25 µL assembled RNP mixture (1 µl SpCas9 2NLS Nuclease (Synthego) (20 pmol/µl, 6 µl target-specific sgRNAs (30 pmol/µl) and 18 µl Opti-MEM^TM^). Controls included a positive control by transfection with sgRNA targeting the previously validated *TRAC* gene (Human TRAC Positive Control, Synthego), and a negative (mock) control. After electroporation organoids were resuspended in an OCM medium, pelleted and singularly seeded in Cultrex®.

Three weeks after electroporation, genomic DNA was extracted using the AllPrep DNA/RNA/protein Kit (QIAGEN). and the targeted locus was amplified. Gel electrophoresis on 2 % agarose gels with SYBR Safe DNA Gel Stain (Invitrogen) visualized the products alongside pBR328 Mix I (Carl Roth) as a DNA marker. A purified PCR product was sent for Sanger sequencing to confirm the edit. Monoclonal organoid lines have been generated by singularization of the pooled cultures, clones picking, and expansion of the individual organoid lines.

#### Western Blot analysis

For the preparation of the total protein samples, organoids were harvested and washed with cold PBS. The pellet was resuspended in 4x Laemmli buffer (Bio-Rad), volume adjusted with PBS to ensure 1x Laemlli concentration, and lysates were heated for 10 min at 95°C. After SDS-PAGE, proteins were transferred to 0.2 µM PVDF membrane by semi-dry Trans-Blot Turbo Transfer System (Bio-Rad), followed by blocking in 3% milk powder and 1.5% BSA blocking solution. After overnight incubation with the primary antibody at 4°C (Table S10) and 3 washing steps in TBS/Tween0.05%, the membrane was incubated with the respective secondary HRP-conjugated IgG-antibody (Table S10) for 1 h at room temperature. After washing, proteins were detected by use of the Cytiva Amersham™ ECL™ Prime Western-Blot-Detection Reagent (Thermo Scientific^TM^) on the ChemiDoc MP Imaging System (Bio-Rad). ß-Actin (Sigma, A5441) Histone-2B (ab1790), or Histone-H3 (ab8580) antibodies were used as endogenous control for the normalization of protein loading.

#### Lentiviral transduction

Ready-to-use lentiviral particles carrying FLAG (DDK)-Myc -agged full-length KRT17, or control vectors (RC201619L1V and PS100064V) have been transduced following the established protocol^55^.In short cell culture plates (24-well format) were pre-coated with a 1:1 mixture of Cultrex and ADF medium and left to solidify overnight at 37⁰C. Organoids were singularized and counted to calculate MOI. An appropriate number of lentiviral KRT17 particles and a control vector were added to the suspension with the addition of polybrene to facilitate the transduction, mixture was transferred to precoded plates and incubated 24h at 37⁰C. The next day cells were retrieved and washed and 3D seeding was performed, according to the standard organoid cultivation protocol. Successful transduction and protein expression of the tagged KRT17 construct has been validated by Western Blotting.

#### RNA Sequencing and Data Analysis

Preparation of the library and NGS Sequencing of the RNA samples from carboplatin-treated and *non-*treated ovarian cancer organoids, isolated by mRNA Easy (Qiagen) kit according to the ‘manufacturer’s protocol, was performed by Novogene Co. Ltd. China, on the Illumina PE150 Novaseq platform, fulfilling stringent quality control criteria (Q30 >95%). The reads were aligned against the index file built from the GRCh38 primary assembly genome (GENCODE release 45) and quantified using the Rsubread package (2.20.0), with the GRCh38 GENCODE v38 GTF file as the reference. Differential gene expression analysis was performed using DESeq2 (version 1.46.0) in the R environment (version 4.4.2). RStudio Packages DESeq2 (1.46.0), ggrepel (0.9.6), dplyr (1.1.4), pheatmap (1.0.12), clusterProfiler (4.14.4), org.Hs.eg.db (3.20.0), AnnotationDbi (1.68.0), DOSE (4.0.0), enrichplot (1.26.05), ggplot2 (3.5.1) were used for the analysis and visualization of differential gene expression data.

#### Mass Spectrometry Analysis

Cell pellets were lysed in Eppendorf tubes with 50 μl of boiling lysis buffer (5% SDS, 100 mM Tris pH 8.5, 5 mM TCEP, and 10 mM CAA). The lysates were incubated at 95 °C for 10 min with mixing (500 rpm) and quantified with Pierce BCA Protein Assay Kit (Thermo Scientific). Protein digestion was performed using a Protein Aggregation Capture (PAC) ^56^-based digestion MagReSyn Hydroxyl beads (ReSyn Biosciences) on a King Fisher Flex system scientific (Thermo scientific).

Samples were mixed with beads were added to the samples at a protein beads ratio of 1:2. For each sample, 200 μl of digestion solution (50 mM triethylammonium bicarbonate) containing Lys-C and Trypsin at an enzyme-to-substrate ratio of 1:500 and 1:250, respectively. Digestion was performed overnight at 37 °C. Protease activity was quenched by acidification with TFA to a final volume percentage of 1% ^57^.

For proteome experiments, 500 ng of digested peptide were loaded in an Evotip Pure (Evosep) following manufacturing instructions. Samples were analyzed using an Evosep column (PepSep EV-1109, 8 cm −150 µm - C18 1.5 µm) and a fused silica (EV-1087 - 20 µm) using an EASY-Spray™ source and interfaced with the Orbitrap Astral Mass Spectrometer (Thermo Scientific). In all samples, spray voltage was set to 1.8 kV, funnel RF level at 40, and heated capillary temperature at 275 °C. Samples were separated on an Evosep One LC system using the pre-programmed gradient for 60 samples per day (SPD – 21 minutes gradient). All experiments were acquired using a data-independent acquisition (DIA) method, cycle time of 0.6 seconds, in a range of 480-1080 m/z, and a maximum injection time (IT) of 30 ms.

#### Data analysis Proteomics

Raw files were analyzed in Spectronaut v19 (Biognosys) with a spectral library-free approach (directDIA+) using the human protein reference 232 database (Uniprot 2022 release, 20,588 sequences) and 233 complemented with common contaminants (246 sequences). Carbamidomethylation of cysteine was set as a fixed modification, whereas oxidation of methionine, N-terminal protein acetylation was set as variable modifications. The enzyme/cleavage rule was set to Trypsin/P, the digest type to specific, and a maximum number of two missed cleavages per peptide was allowed. The maximum number of variable modifications per peptide was set to 5 and method evaluation was turned off.

The data analysis of proteome data was performed in R version 4.4.0 (R Core Team (2021), R foundation for statistical computing) with R Studio 2024.4.0.735. The output files were analyzed using Rstudio with the following packages, dplyr (1.1.4), tidyverse, (2.0.0), ggrepel (0.9.6), Limma (3.60.6), EnhancedVolcano (1.22.0), clusterProfiler (4.12.6), enrichplot (1.24.4), org.Hs.eg.db (‘3.19.1).

#### Fixation and embedding of organoids

Fixation of organoids in Epredia™ Richard-Allan Scientific™ HistoGel™ (Thermo Scientific™) for subsequent paraffin embedding, micro-sectioning, and staining procedures followed our institutional protocol ^12^. In summary, organoids were released from the Cultrex® BME 2 matrix by cold ADF++, fixed in 4% Paraformaldehyde (PFA), and washed twice in PBS. Fixed organoids were gently resuspended in 100 µl of pre-heated HistoGel™, and the solidified HistoGel™ droplet was processed to paraffin embedding and slicing.

#### Immunofluorescence staining of organoids and tissue

Immunofluorescence staining was performed as previously described ^12^. In short, slides were deparaffinized in Roticlear® (Carl Roth), followed by rehydration in a descending ethanol series, antigen retrieval by heating in a steamer to 100°C with TRIS/EDTA-solution and permeabilization for 15 min in 1% Triton® X-100 (in PBS) solution (Sigma-Aldrich®). After blocking with 10 % serum, and primary antibody incubation at 4°C for at least 16 h, slides were stained with secondary IgG antibodies for 2 h at room temperature. DAPI counterstaining (Thermo Scientific™) visualized nuclei. Mowiol® 4-88 Water soluble Mounting Medium (Carl Roth) sealed the slides for imaging on a Leica SP8 confocal microscope.

#### Immunohistochemistry of TMA sections

After the removal of paraffine (Roticlear, and decreasing ethanol range), slides were incubated in antigen-retrieval solution pH 8,0 (Novocastra RE7116), for 2x 15 min in the heated Microwave and left to cool down for 20 min at room temperature. Next, samples were blocked for 10 min with Protein Block (Agilent Technologies, X0909). Primary antibody anti KRT17 (clone SPM560, ab 233912), diluted 1:250, was incubated for 1h at room temperature. After washing, (PH 7,5 TRIS buffer), samples were treated with AP Polymer anti-Mouse (Zytomed systems, ZUC-077) for 30 min, washed and Vector Red substrate was added for 20 min (Vector, SK-5100). Excess dye was washed out for 10 min under running water, counterstaining with Hematoxylin (Vector, H-3401) was performed for 10 sec and the washing step was repeated. Samples we mounted with Aquatex (Merck, 1.08562.0050).

#### TMA Staining evaluation

The ovarian cancer TMA slides were visualized by the automated Keyence Microscope BZ-X800 at a 20x magnitude with automated image stitching. The QuPath Bioimage Analysis Software ^38^ was used for IHC analysis of KRT17 expression on TMA slides. An automated script ensured accurate, consistent, and objective IHC scoring (Table S11). Stains were separated by color deconvolution after smoothing. QuPath’s Pixel and Object Classifier algorithms were interactively trained on more than 30 selected heterogeneous training images within the study collective to automatically annotate regions of interest and to distinguish epithelial tumor cells from all other detections. The QuPath’s Positive Cell Detection algorithm was applied for cell segmentation and counting. Intensity thresholds were determined for either negative, weakly positive, or strongly positive cells based on the cell KRT17 optical density sum (negative <0.09, weakly positive 0.09-0.17, strongly positive >0.17, Fig. 4D). Automated batch analysis was applied to all 384 duplicated TMA samples, providing data for further statistical analysis.

#### Statistical analysis

For the TMA analysis, the total cell count for tumor and stroma cells and positive cell detection were pooled for both sample duplicates before further statistical analysis. A tailored immunoreactivity score was established. For this purpose, weakly positive cell count was added to the twofold count of strongly positive cells, then divided by the total tumor cell count and multiplied by 100, resulting in a range of 0 (all tumor cells negative for KRT17) to 200 (all tumor cells strongly positive for KRT17). We named this adapted immunoreactivity score “K-score”. Samples with less than 1150 detected tumor cells in total were excluded from further statistical analysis, based on the required sample size to classify the sample based on a 5% cutoff with >99,99% confidence. Clinical information, including data about survival and disease progression, was related to the immunohistochemical scoring by descriptive statistical analysis, Kaplan-Meier analysis, and Multivariate Cox regression analysis by the IBM SPSS Statistical Software (version 29.0.1.0 (171)) as well as Survival (version 3.7.0) and survminer (0.5.0) packages in R Data analysis and visualization by the web application *Cutoff Finder* ^46^ facilitated the determination of an optimal cutoff point for patient stratification.

#### Data Visualization

Data visualization was performed using GraphPad Prism Software Version 10.1.1. software was used to generate concentration-response curve diagrams for organoid counting and mRNA expression, analyze cell viability data, and create graphical representations of the experimental results obtained in Qupath (organoid count and surface area data). R packages survival, surminer, dplyr, foreign, and ggplot2 were used for data processing and generating plots of survival analysis, and HR plots of Cox proportional model analysis.

